# Beyond Segmentation Accuracy: Evaluating Biological Fidelity in Spatial Pathology

**DOI:** 10.64898/2026.10.17.759841

**Authors:** Nevenka Dimitrova, Marjan Stoimchev, Aveline Filliol, Sara Flowers, Anthony Cipriano, Pranav Suresh, Paul Romesser, Scott Lowe

## Abstract

Multiplex immunofluorescence (MxIF) resolves cellular identity, state, and spatial organization within tumor tissue, enabling higher-order representations of cancer ecosystems including immune infiltration, multicellular neighborhoods, and tissue niches. Yet these analyses depend on a deceptively simple first step: accurately identifying individual cells. Segmentation errors can alter marker quantification, cellular phenotypes, cell–cell relationships, and ultimately the spatial organization inferred from an image. Consequently, improvements in conventional segmentation benchmarks matter biologically only if they propagate to more faithful representations of the tumor microenvironment.

We test this relationship directly using an expert-curated benchmark of 31,924 expert-reviewed cells on three slides under a leave-slide-out protocol. We compare a domain-specific self-supervised encoder, adapted to multiplex tissue imaging, with supervised, microscopy-specific, prompted generalist, and general-purpose vision models, and pass every method’s predicted cells through an identical downstream pipeline. We evaluate fidelity across four levels: cell segmentation, phenotype assignment, spatial neighborhood structure, and tissue organization, comprising niches and tumor–immune interactions.

The resulting hierarchy reveals that these levels do not improve in lockstep. Cellular phenotype assignment is comparatively robust to segmentation differences, whereas missed cells increasingly disrupt neighborhood structure and tumor–immune interactions. As a stringent spatial-interaction test, tumor–CD8^+^ contact recovery is poorly predicted by conventional segmentation quality (Panoptic Quality: *ρ* = 0.18), while errors specifically at tumor–immune interfaces are strongly associated with contact fidelity (*ρ* = −0.80). Methods with nearly identical segmentation accuracy can therefore produce substantially different biological representations. These results argue that cell segmentation for spatial pathology should be evaluated not only by boundary and instance accuracy, but also by preservation of the biological relationships that downstream analyses are intended to measure.

## 1 Introduction

Multiplex immunofluorescence gives a spatially resolved view of cancer tissue in which individual cells carry morphology, molecular phenotype and local context at once. This supports analyses well beyond cell counting: immune infiltration patterns, tumor–immune interactions, multicellular neighborhoods and tissue niches, representations increasingly used to characterize tumor ecosystems and, ultimately, to stratify patients.

These analyses inherit the errors of the first computational step. A segmentation error alters the pixels assigned to a cell, and with them its marker profile, its phenotype, its spatial relationships and the niche assigned to a region; in dense tumor tissue a merge between an immune and a tumor cell can matter more than a boundary displacement costing the same in overlap metrics. Segmentation is therefore not only a computer-vision benchmark but the start of a chain of scientific inference.

We therefore ask what it means for a foundation model to be better for spatial pathology: whether gains measured at the cell-segmentation level preserve the biological information required downstream. Such models should be judged by what their representations preserve, not by an intermediate computer-vision score alone, so we organize the evaluation around four progressively more biologically meaningful levels (Table 3), tracing errors from cell boundaries through phenotypes and neighborhoods to niches and tumor–immune organization, with KODIAK [29], an existing self-supervised model for MxIF, as a controlled case study. The hierarchy also tests whether segmentation errors are biologically exchangeable: errors of equal geometric magnitude may have very different consequences depending on which cells they affect and where.

### Contributions

We make three contributions: (1) a controlled framework for evaluating cell segmentation across a hierarchy from instance accuracy to biological representation; (2) a multi-model MxIF benchmark showing that segmentation, phenotype, neighborhood, niche, and tumor–immune fidelity do not improve in lockstep; and (3) evidence that biologically localized segmentation errors at tumor–immune interfaces predict downstream contact fidelity substantially better than conventional global segmentation metrics.

## 2 Related Work

Cell segmentation in fluorescence microscopy is dominated by flow- and distance-regression models trained on large annotated corpora [2, 3, 5, 6, 7, 8], by commercial platforms such as HALO [9], still the standard in pathology laboratories, and by promptable generalist models of the SAM family [10, 11, 12], whose prompts remove training altogether; we place each family under one review-aware, region-level protocol. Self-supervised backbones of the DINO lineage [13, 14, 15] give strong frozen features for dense prediction, and pathology foundation models continue their pretraining on histology at scale [20, 21, 22], but whether that helps instance-level tasks over the foundation they start from is largely untested; our comparison varies only the encoder initialization. Downstream, spatial single-cell analyses of multiplexed tissue [23, 24] gate markers into phenotypes, build cell graphs and cluster neighborhood compositions into niches whose organization correlates with outcome [25, 26, 27]; every step consumes cell boundaries.

## 3 Benchmark Dataset and Evaluation Protocol

### 3.1 Slides, annotations, and quality review

The benchmark is built on three CellDIVE [23] MxIF whole-slide images of mouse colorectal cancer tissue acquired with the same nine-marker panel, the nuclear counterstain DAPI, the nuclear-envelope marker LAP2, the epithelial marker PanCK, the immune markers CD45, CD3, CD4, CD8 and FOXP3, and uPAR, at 0.325 *µ*m per pixel (Figure 1, Table 1). We refer to them as Slide 1, Slide 2 and Slide 3; their roles under the protocol of Section 3.2 are fixed in advance, and no number uses a slide in any other role.

**Table 1.** The benchmark data: every count used in this paper. Cells are HALO cell objects owned by an annotated region and *confirmed* or *flagged* by the review, or lying *outside* the review (Slide 3’s unreviewed field, and four Slide 1 contours without a review row); tiles are 256^2^ windows on a 192-pixel grid where labeled and a 128-pixel grid where not. Slide 1 measures 20,437*×* 13,048 pixels, Slide 2 18,626*×* 14,977 and Slide 3’s annotated scan region 16,839*×* 35,071, all at 0.325 *µ*m per pixel. Phenotype labels: Table 4.

| Slide, stage | Regions |  | Cells |  |  |  | Tiles |  |
| --- | --- | --- | --- | --- | --- | --- | --- | --- |
|  | all | reviewed | all | confirmed | flagged | outside | labeled | unlabeled |
| **Slide 1** · development | 16 | 16 | 21,657 | 16,985 | 4,668 | 4 | 628 | 4,478 |
| training | 14 | 14 | 19,259 | 15,063 | 4,192 | 4 | 550 | 4,478 |
| model selection (R14, R15) | 2 | 2 | 2,398 | 1,922 | 476 | 0 | 78 | — |
| **Slide 2** · test (R1–R6) | 6 | 6 | 7,662 | 6,152 | 1,510 | 0 | 259 | 4,103 |
| **Slide 3** · second test | 4 | 3 | 3,451 | 1,672 | 937 | 842 | 72 | — |

**Figure 1:**
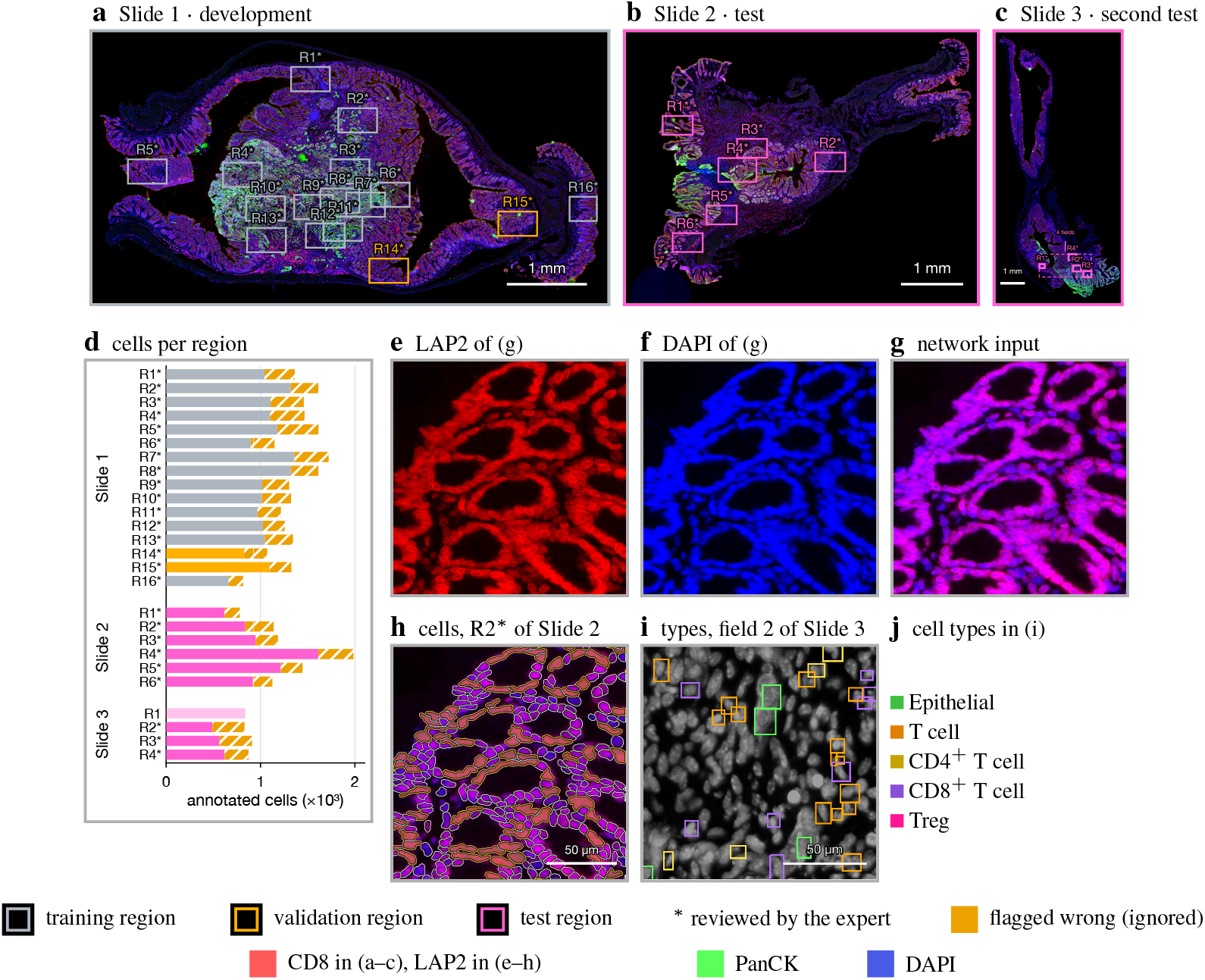
The benchmark. (a–c) The three slides in CD8, PanCK and DAPI, exposed 1.7 *×* harder than Figure 6a for a panel this small, with annotated regions outlined by their role and reviewed regions starred (1 mm scale bars). (d) Annotated cells per region: solid, confirmed by the review; pale, the one unreviewed field of Slide 3, scored nowhere; hatched amber, flagged. (e–g) A 512^2^ crop (166 *µ*m) of test region R2 of Slide 2: the two channels and the input they form. (h) The same input with the expert annotation. (i) A 140 *µ*m crop of field 2 of Slide 3, every labelled cell drawn at its deepest class (the classes nest); a type label locates a cell, it never defines a boundary. *Colour:* DAPI is blue in every panel of the paper, as the network input itself puts it; (e–h) add LAP2 on red, (a–c) CD8 on red and PanCK on green. No pixel values are altered anywhere, and every panel is windowed on its own slide’s normalization.

Rectangular regions were segmented by the HALO platform [9] into closed cell contours: DAPI-anchored objects drawn from the DAPI and LAP2 channels with a small cytoplasmic margin (median equivalent diameter 7.5 *µ*m on Slide 1), which we call cells throughout. An expert then reviewed every region cell by cell, inspecting every object and flagging the incorrect ones, predominantly merges, as *wrong*. The review covers all sixteen regions of Slide 1, all six of Slide 2 and three of Slide 3’s four fields, 31,924 cells in all (Table 1), and every scored cell of this paper comes from them. These flags define an ignore mask honored everywhere: flagged cells leave training losses and every reported metric, and a prediction on one is neither a hit nor a false positive, so neither training nor evaluation rewards known annotation errors. The four Slide 1 contours without a review row enter training with their contours as provided, and the one unreviewed field of Slide 3 is left out entirely; no number in this paper is computed on them.

#### Phenotype annotations

Three sets of expert phenotype labels serve the phenotyping level (Section 4.3) and never touch segmentation training or scoring (Table 4): on Slide 1, one 1,024^2^-pixel window inside training region R1, tagged exhaustively per session so untagged cells are negatives, which calibrates the gates; on Slide 2, four such windows, held out with the slide; on Slide 3, bounding boxes in two reviewed fields for CD3^+^ T cells with their CD4^+^ and CD8^+^ subsets, FOXP3^+^ regulatory T cells and PanCK^+^ cells. Slides 2 and 3 are scored once each; no model saw either at any stage.

### 3.2 Leave-slide-out protocol

Every design decision that could adapt a model to the evaluation data is confined to Slide 1, and Slide 2 is accessed only for the final evaluation of each completed run. Slide 1 serves as the *development slide*: KODIAK is continually pretrained on its tissue tiles (Section 4.1); every supervised model is trained on fourteen of its regions, all reviewed so that the ignore mask is active throughout training, and selected on regions R14 and R15 at the epoch of minimum validation loss (Table 1); and the remaining degrees of freedom, namely optimization hyperparameters, decoding thresholds, phenotype gates and analysis settings, are established there. Slide 2 is the *test slide*: none of its pixels enters pretraining, training or model selection, and it is scored once per model on whole regions (Section 3.3). The validation regions share no pixel with a training region (annotated regions of Slide 1 partially overlap). A fixed manifest records the exact tiles, augmentation draws and ordering, so every trained model consumes the identical stream; each is trained three times under independent random initializations, which also reorder its tiles and redraw its augmentations. Every number we report for a method is the mean over its three runs. Run-to-run variation is small enough at the precision we report that we do not carry it through the tables, and differences between methods are assessed instead by paired per-region comparisons. The numbers therefore measure generalization to unseen slides: Slide 2 and Slide 3 are both scored this way, each once, after every model was frozen.

### 3.3 Region-level evaluation

We report all results at the level of whole annotated regions of the test slide, each contributing one complete ground-truth map and one complete prediction map. Predictions come from 256^2^ tiles on the training grid, stitched by an ownership rule (an object is taken whole from the tile whose core contains its centroid) so no object is cut by a seam, and matched by Hungarian assignment [31] on instance IoU. We report F1 and average precision at IoU 0.5, AP over IoU 0.5–0.9 (AP_.5:.9_), Panoptic Quality [PQ; 30], precision, recall, and the count ratio (predicted over true instances; 1 is perfect calibration). Because every test region is fully reviewed, every scored cell has been individually confirmed. Each reference cell is also classified into an error taxonomy (correct, missed, merged, split or poor boundary) and each prediction touching no reference cell as gained, with paired per-region differences under bootstrap intervals (Appendix A.4.1).

## 4 Methods

### 4.1 Domain-specific pretraining

KODIAK is a published self-supervised objective for multiplex tissue imaging [29]: a ViT/16 student [16] trained against an exponential-moving-average teacher by masked prediction and cross-view alignment over a learned prototype codebook. The encoder benchmarked here is initialized from DINOv3 [15] and continually pretrained under that objective on Slide 1 alone, over its training-split tiles (labels never read), at ViT-B/16 to match Cellpose’s encoder. Pretraining adds one phenotype marker, sampled per sample, on the green channel, so the representation sees lineage context that the segmentation input never carries. The identical network initialized from DINOv3 without continued pretraining is the control in every comparison, so the difference isolates what continued in-domain pretraining contributes at slide scale. No external MxIF corpus enters any number reported here.

### 4.2 Segmentation methods

We compare KODIAK with representatives of the model families in current use, spanning four levels of supervision (the blocks of Table 2), all evaluated under the protocol of Section 3.3 on the same test regions with the same ignore mask.

**Table 2.**
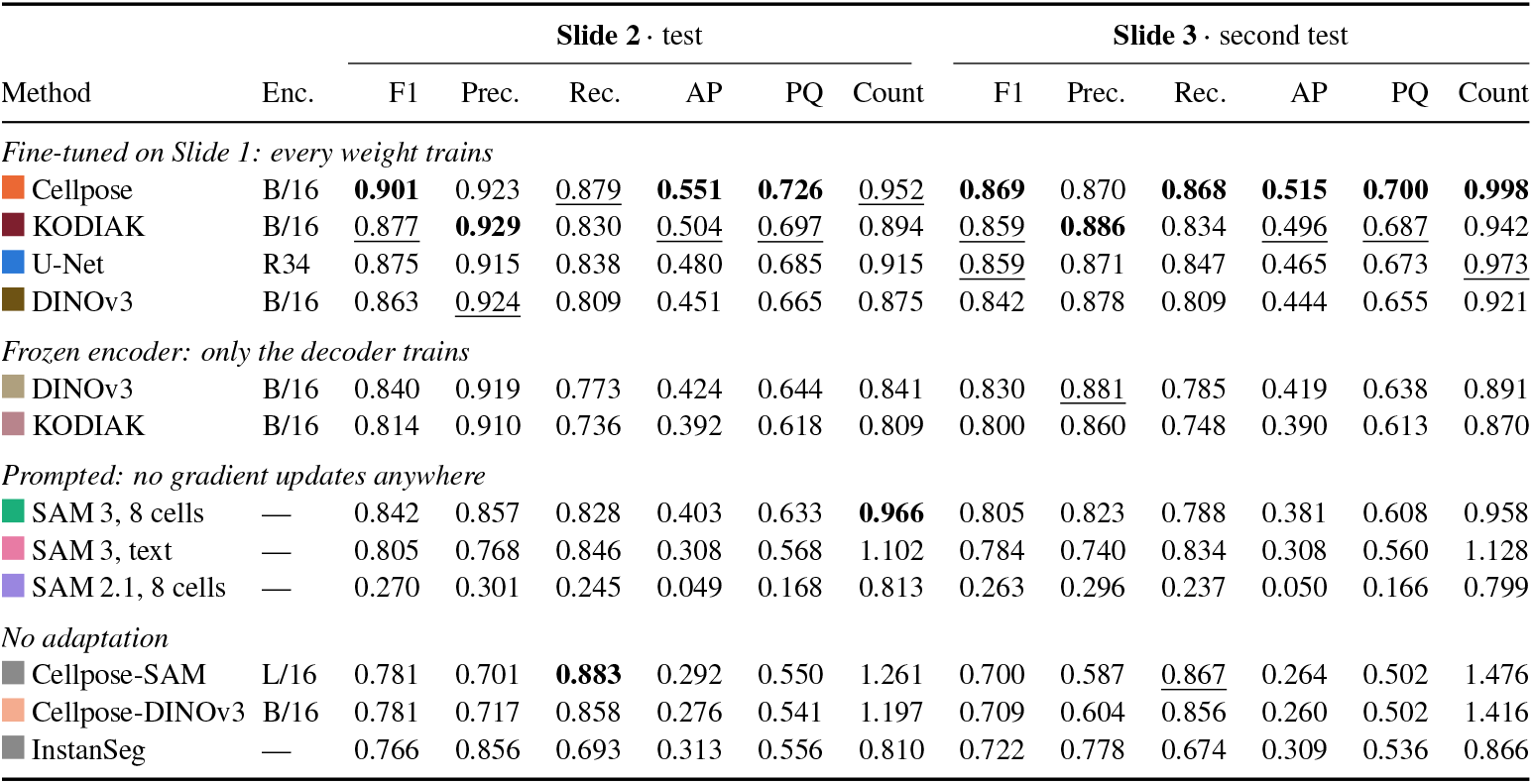
Region-level segmentation on the two held-out slides. Slide 2: every method is scored on the identical set, all six regions, the same 6,152 confirmed cells, the same 1,510 excluded by the reviewer’s flag. Slide 3: its three reviewed fields, 1,672 confirmed cells, scored once after every model was frozen; it contributes no training or selection signal anywhere. *Enc*. is the encoder; the block headings give the supervision each group received. Rows are ordered by Slide 2 F1 within each block; **best** and <u>second best</u> per column, Count closest to 1. AP is averaged over IoU .5–.9; AP at .5 equals F1*/*(2 − F1) and is omitted. Every trained row is the mean over three independent training runs (Section 3.2).

|  |  | Slide 2 · test |  |  |  |  |  | Slide 3 · second test |  |  |  |  |  |
| --- | --- | --- | --- | --- | --- | --- | --- | --- | --- | --- | --- | --- | --- |
| Method | Enc. | F1 | Prec. | Rec. | AP | PQ | Count | F1 | Prec. | Rec. | AP | PQ | Count |
| Fine-tuned on Slide 1: every weight trains |  |  |  |  |  |  |  |  |  |  |  |  |  |
| Cellpose | B/16 | **0.901** | 0.923 | <u>0.879</u> | **0.551** | **0.726** | <u>0.952</u> | **0.869** | 0.870 | **0.868** | **0.515** | **0.700** | **0.998** |
| KODIAK | B/16 | <u>0.877</u> | **0.929** | 0.830 | <u>0.504</u> | <u>0.697</u> | 0.894 | <u>0.859</u> | **0.886** | 0.834 | <u>0.496</u> | <u>0.687</u> | 0.942 |
| U-Net | R34 | 0.875 | 0.915 | 0.838 | 0.480 | 0.685 | 0.915 | <u>0.859</u> | 0.871 | 0.847 | 0.465 | 0.673 | <u>0.973</u> |
| DINOv3 | B/16 | 0.863 | <u>0.924</u> | 0.809 | 0.451 | 0.665 | 0.875 | 0.842 | 0.878 | 0.809 | 0.444 | 0.655 | 0.921 |
| Frozen encoder: only the decoder trains |  |  |  |  |  |  |  |  |  |  |  |  |  |
| DINOv3 | B/16 | 0.840 | 0.919 | 0.773 | 0.424 | 0.644 | 0.841 | 0.830 | <u>0.881</u> | 0.785 | 0.419 | 0.638 | 0.891 |
| KODIAK | B/16 | 0.814 | 0.910 | 0.736 | 0.392 | 0.618 | 0.809 | 0.800 | <u>0.860</u> | 0.748 | 0.390 | 0.613 | 0.870 |
| Prompted: no gradient updates anywhere |  |  |  |  |  |  |  |  |  |  |  |  |  |
| SAM 3, 8 cells | — | 0.842 | 0.857 | 0.828 | 0.403 | 0.633 | **0.966** | 0.805 | 0.823 | 0.788 | 0.381 | 0.608 | 0.958 |
| SAM 3, text | — | 0.805 | 0.768 | 0.846 | 0.308 | 0.568 | 1.102 | 0.784 | 0.740 | 0.834 | 0.308 | 0.560 | 1.128 |
| SAM 2.1, 8 cells | — | 0.270 | 0.301 | 0.245 | 0.049 | 0.168 | 0.813 | 0.263 | 0.296 | 0.237 | 0.050 | 0.166 | 0.799 |
| No adaptation |  |  |  |  |  |  |  |  |  |  |  |  |  |
| Cellpose-SAM | L/16 | 0.781 | 0.701 | **0.883** | 0.292 | 0.550 | 1.261 | 0.700 | 0.587 | <u>0.867</u> | 0.264 | 0.502 | 1.476 |
| Cellpose-DINOv3 | B/16 | 0.781 | 0.717 | 0.858 | 0.276 | 0.541 | 1.197 | 0.709 | 0.604 | 0.856 | 0.260 | 0.502 | 1.416 |
| InstanSeg | — | 0.766 | 0.856 | 0.693 | 0.313 | 0.556 | 0.810 | 0.722 | 0.778 | 0.674 | 0.309 | 0.536 | 0.866 |

#### Baselines

The U-Net baseline is an ImageNet-initialized ResNet-34 encoder [18] under a U-Net decoder [1]. KODIAK and DINOv3 share one segmentation network and differ only in the initialization of its ViT encoder (the MxIF-pretrained teacher of Section 4.1 or the released DINOv3 weights), so that their comparison isolates the contribution of continued pretraining. Cellpose [2, 4] is the released flow-field model with a DINOv3 ViT-B/16 backbone (86M parameters), evaluated fine-tuned on Slide 1 and without adaptation; the released Cellpose-SAM (ViT-L/16) and InstanSeg [8] are evaluated as distributed. SAM 3 [12] and SAM 2.1 [11] receive no gradient updates: SAM 3 is prompted with the boxes of eight annotated cells drawn deterministically from the training regions, or with the text prompt “cell”; SAM 2.1, having no exemplar interface, with a prototype embedding over the same eight (Appendix A.3.2).

#### Input

Every model receives the same two-channel image (Figure 1e–g): LAP2 on red and DAPI on blue, the channels from which the reference cells were drawn, with green left empty. Each marker is scaled to [0, 1] by slide percentiles computed without labels, then by the ImageNet [19] normalization the pretrained encoders expect (Appendix A.3.1). The prompted and zero-shot models see that same image, with exemplars cut from the same rendering, so they differ from the trained rows in never having been fitted to it, not in what they are shown.

#### Training

All slide-trained models regress the Cellpose target triplet (a two-channel flow field toward each cell center and a cell-probability logit) under a loss masked by the expert ignore mask, and instances are decoded by Cellpose’s flow-following procedure. Every model gets the identical recipe and tile stream (Appendix A.3.1); the epoch of minimum validation loss on R14 and R15 is kept, and each model is evaluated on the test slide once.

### 4.3 Cell phenotyping and spatial analysis pipeline

To isolate the effect of segmentation, everything downstream is held fixed: every method’s cells, and the expert’s reference cells, pass through the identical pipeline, so any difference at a higher level is attributable to the cell boundaries alone. A cell’s marker value is the mean intensity inside its mask, in the slide-normalized units of Section 4.2. Phenotypes follow an expert-provided hierarchical taxonomy (epithelial by PanCK, immune by CD45, T cells by CD3 subdivided by CD4 and CD8, regulatory T cells by FOXP3, and the uPAR^+^ epithelial state), one threshold per marker, applied to expert and predicted boundaries alike and calibrated by cross-validation to the human labels on the development slide only. Spatial neighborhoods are *k*-nearest-neighbor graphs (*k* = 6) over cell centroids; cellular niches are *k*-means clusters of ten-cell neighborhood compositions [25, 26], derived once from the expert’s cells and assigned to every method’s cells through its own graph and phenotypes.

## 5 Results

The results follow the hierarchy of Table 3, every method’s cells passing through the identical pipeline of Section 4.3. Section 5.1 reports the segmentation level and Sections 5.2–5.5 the three levels above it, for every method; the prose of the downstream sections follows the six slide-trained rows, and the released and prompted rows are discussed where they differ.

**Table 3.** Evaluation hierarchy linking technical segmentation performance to increasingly biologically meaningful representations of the tumor microenvironment.

| Level | Question | Interpretation |
| --- | --- | --- |
| 1. Cell segmentation | Did we recover individual cells and their boundaries correctly? | Technical benchmark |
| 2. Cell phenotype | Did segmentation preserve cellular identity and marker measurements? | Cellular representation |
| 3. Neighborhoods | Did we preserve spatial relationships and local cellular composition? | Spatial representation |
| 4. Tissue organization | Do we recover the same niches and tumor–immune interactions? | Biologically meaningful representation |

**Table 4.** The human phenotype labels of the phenotyping level (Section 4.3). They never touch segmentation training or scoring.

| Phenotype labels | Slide | Location | Cells | Labels |
| --- | --- | --- | --- | --- |
| tags, annotation tool | 1 | 1,024<sup>2</sup>-pixel window in R1 | 1,283 | PanCK<sup>+</sup> 1,006; T cell 40; CD4<sup>+</sup> T cell 26; CD8<sup>+</sup> T cell 12; untagged cells are negatives |
| tags, annotation tool | 2 | four 1,024<sup>2</sup>-pixel windows | 2,468 | PanCK<sup>+</sup> 1,982; T cell 188; CD4<sup>+</sup> T cell 147; CD8<sup>+</sup> T cell 71; Treg 80; held out with the slide |
| bounding boxes | 3 | reviewed fields 2 and 4 | 417 | 756 boxes. Field 4: CD3<sup>+</sup> 97, CD3<sup>+</sup>CD4<sup>+</sup> 72, CD3<sup>+</sup>CD8<sup>+</sup> 14, FOXP3<sup>+</sup> 13, PanCK<sup>+</sup> 14; field 2: CD3<sup>+</sup> 96, CD3<sup>+</sup>CD4<sup>+</sup> 51, CD3<sup>+</sup>CD8<sup>+</sup> 89, FOXP3<sup>+</sup> 13, PanCK<sup>+</sup> 297; and 76 further CD3<sup>+</sup> boxes in an area with no segmentation |

### 5.1 Domain-specific pretraining improves fine-tuned MxIF cell segmentation

Table 2 reports segmentation accuracy on the six test regions of Slide 2. Since its means average over regions of different size and difficulty, we assess each comparison by the paired per-region difference in F1, averaged over runs, with a bootstrap 95 % confidence interval over the six regions.

#### Model comparison

Among the models trained on Slide 1, Cellpose leads at F1 0.901, above every other model on every region (+0.024 over KODIAK, CI +0.020 to +0.028), and the fine-tuned KODIAK and the U-Net are indistinguishable (0.877 and 0.875; CI − 0.003 to +0.003), both above the DINOv3 control (0.863). Precision is nearly uniform across the trained rows (0.91–0.93), so the differences are differences in recall, from 0.736 to 0.879. Continued MxIF pretraining points one way when the encoder is fine-tuned and the other when it is frozen: fine-tuned, KODIAK beats its DINOv3 control on *every* test region, by +0.014 (CI +0.011 to +0.018), through recall (+0.021) at matched precision; frozen, the order reverses just as consistently (−0.024, CI −0.030 to −0.018, again on all six regions). The gain is in adaptability, not decodability. What separates Cellpose from the same DINOv3 backbone fine-tuned here (0.863) is therefore supervision rather than pedigree: its cell-corpus pretraining is worth +0.038, and it shows almost entirely in recall (+0.048 over KODIAK at −0.005 precision).

#### Prompted models are competitive with little supervision

SAM 3 prompted with the bounding boxes of eight annotated cells reaches F1 0.842 with the best count calibration of any method (0.966), level with the frozen DINOv3 probe (CI −0.005 to +0.013), above the frozen KODIAK probe on every region, and within 0.033 of the U-Net trained on 550 annotated tiles (CI −0.043 to −0.024). The text prompt “cell”, with no visual supervision, reaches 0.805, while SAM 2.1 performs poorly when the same eight exemplars are averaged into a prototype embedding (F1 0.270): few-shot performance depends strongly on how exemplars are incorporated. The released models without adaptation, Cellpose-SAM, Cellpose-DINOv3 and InstanSeg, reach 0.77–0.78; the two Cellpose releases recover more cells than several trained rows (recall 0.86–0.88) but predict 1.2–1.3 objects per reference cell.

#### The ordering transfers to a second slide

Scored once on the three reviewed fields of Slide 3 under the identical protocol, every trained model loses a little accuracy — F1 falls by 0.010 to 0.032, a mean of 0.018 — and the ranking is unchanged (Spearman +0.97 between the two slides over the twelve rows). Cellpose still leads (0.869), KODIAK and the U-Net still tie (0.859 each), KODIAK still beats its DINOv3 control fine-tuned (+0.017, every field) and still loses to it frozen (−0.030). Precision falls by about 0.05 while recall holds, so the second slide adds false positives rather than misses, some plausibly cells the reference lacks (Appendix A.4.2). The prompted and released rows degrade more (− 0.037 for SAM 3 with exemplars, −0.07 to − 0.08 for the two Cellpose releases, through precision), which places SAM 3 below every fine-tuned row and the frozen DINOv3 probe on a slide it was not prompted from. Only the segmentation level is measured on Slide 3; the phenotype, neighborhood, niche and tumor–immune analyses that follow are computed on Slide 2 alone.

#### Error analysis

Figure 7 attributes the recall differences. Missed cells dominate (9–18 % of the reference cells for the trained rows; 8 % for Cellpose), followed by merges (3–5 %), with splits and poor boundaries together below 4 % for every trained model. Missed cells are small and crowded: recall is 0.44–0.61 for cells of 15–25 *µ*m^2^ and 0.48–0.77 for cells whose nearest neighbor lies within 5 *µ*m, against at least 0.90 above 50 *µ*m^2^ or 0.92 beyond 9 *µ*m, and Cellpose’s advantage concentrates in exactly these bins. Of the confirmed cells 17.0 % are found by all twelve methods and 2.7 % by none, and 2.3 % by no run at all; whether those are annotation errors that survived review or genuinely hard cells needs an independently re-annotated field (Section 7).

### 5.2 Cell phenotypes are robust to segmentation differences

Marker values and phenotypes are measured inside the expert boundary and inside each method’s matched boundary of every confirmed test cell, and disagreements are stratified by the error class of Section 3.3. Across the six slide-trained rows, lineage agreement is 95.8–96.9 % and full-type agreement 94.3–95.9 %, far narrower than the segmentation spread that produced it; the released and prompted rows sit at 93.4–94.5 % full-type agreement, and only SAM 2.1 falls away (84.1 %). The loss concentrates in one error class: a phenotype survives a correctly matched cell 96.8–97.5 % of the time but only 89.4–91.1 % when the cell has been merged with a neighbor, so a merge is about three times as likely to change a cell’s identity. The ordering here reproduces the segmentation ordering but compresses it heavily: measured on the cells a method recovers, phenotyping is robust to how well it recovered them.

### 5.3 Missed cells disrupt spatial neighborhood structure

Among the cells a method recovers, the *k*-nearest-neighbor graph is nearly intact: edge recall is 0.972–0.976 across the trained rows and essentially method-independent. Scored against the expert’s complete graph it falls to 0.669–0.837 and neighborhood composition drifts by an *L*_1_ of 0.125–0.188, so the damage is attributable to the cells never found rather than to displaced boundaries: a missed cell takes all of its relationships with it, a found cell keeps them even when its boundary is imperfect. Heterotypic contact recall follows the same pattern (0.627–0.824), and the frozen KODIAK probe, which recovers the graph among its matched cells as well as any method (0.975), is last on both whole-graph measures because it finds fewest. The released Cellpose rows, which over-predict, keep the most expert edges (0.92–0.93) at the lowest edge precision.

### 5.4 Niche recovery is niche-dependent

Niches are derived once from the expert cells and assigned to every method’s cells through its own graph and phenotypes. Agreement is 87.2–91.6 % of cells across the trained rows, but recovery is niche-dependent: the epithelial core is recovered on 89.4–95.9 % and the immune-infiltrated niche on 88.7–92.7 %, while the interface niche between them, the one carrying the tumor–immune biology of interest, is recovered on only 76.7–83.8 %, and the largest niche-abundance error grows from 0.4 points for the U-Net to 2.9 for the frozen KODIAK probe. Interface recall separates the methods more than overall niche agreement does, and it is not the segmentation ordering: the U-Net (83.7 %) and DINOv3 (83.8 %) recover it better than KODIAK (80.7 %) and Cellpose (82.6 %).

### 5.5 Conventional segmentation metrics fail to predict tumor–immune contact fidelity

Finally, as a deliberately stringent test of error propagation, we use a spatial interaction endpoint: contact between PanCK^+^ tumor cells and CD8^+^ T cells. This focused case study contains 40 expert-defined contacts across the six test regions (Figure 2 reads one) and requires both cells, their phenotypes, and their spatial relationship to survive. Across the trained rows contact F1 spans 0.45–0.62, about twice the spread of the segmentation F1 of the same runs (0.814–0.901), and it does not preserve their ordering: the U-Net and KODIAK are indistinguishable as segmenters (0.875 against 0.877 F1) yet differ by 0.08 in contact F1 (0.58 against 0.50), and the frozen DINOv3 probe (0.47) comes within 0.03 of KODIAK despite being 0.037 F1 behind it. Across all run–region points, Panoptic Quality correlates with contact F1 at Spearman +0.18 (F1 +0.21, Dice +0.19), while the error rate restricted to tumor–immune interfaces correlates at −0.80 (Figure 4a, b). The mechanism is visible in the accounting (Figure 3a): contacts are lost because one of the two cells was never matched (35–52 % of expert contacts) far more often than because a phenotype flipped (10–14 %) or an adjacency shifted (0–2 %). The expert calls one of the six test regions CD8-infiltrated and five immune-excluded [28]; the methods reproduce that call on five to six regions.

**Figure 2:**
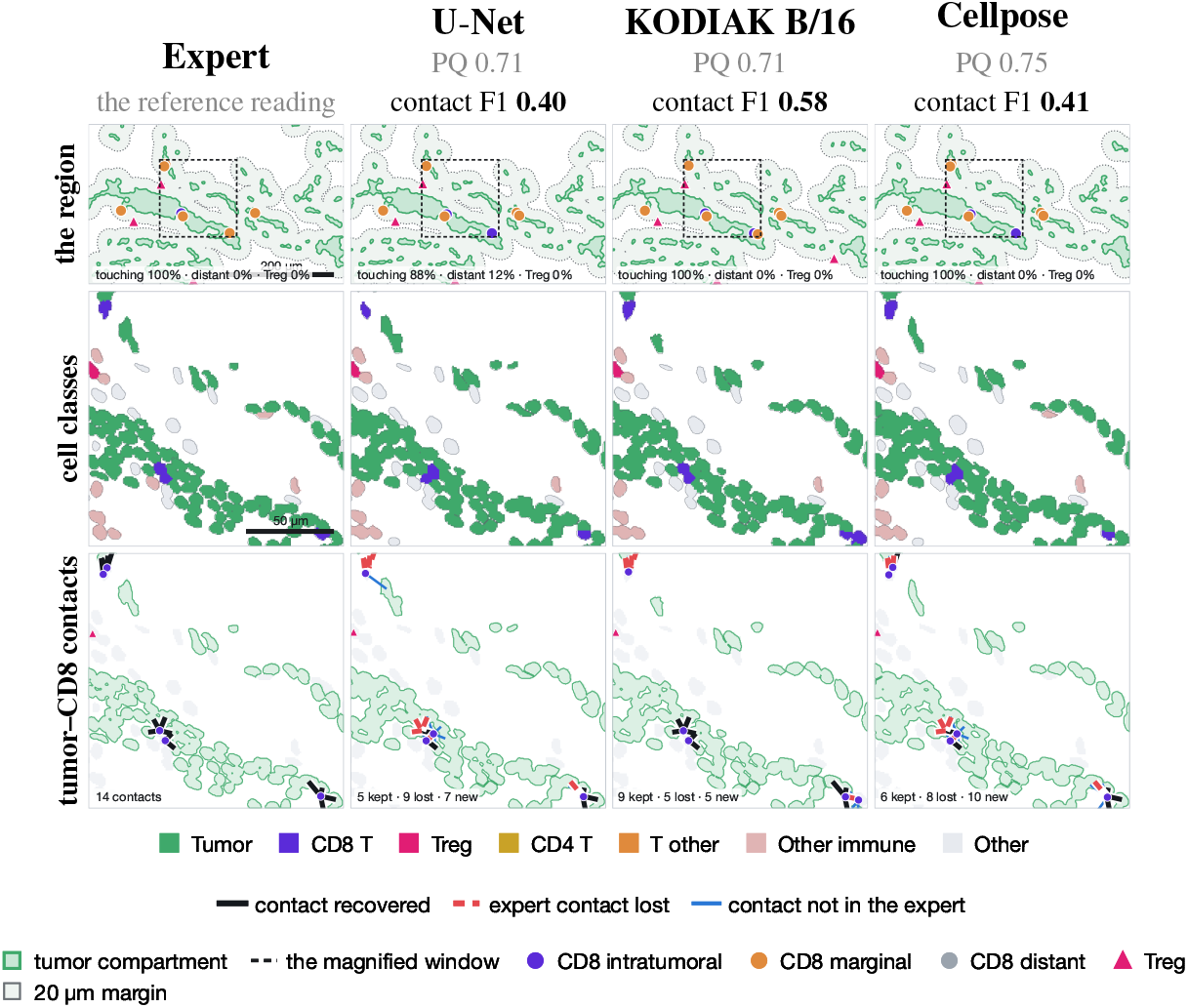
Similar segmentation quality, different biological readings. Test region R1, selected by a rule fixed in advance (Appendix A.5.1). Columns: the expert reference and three methods, one run drawn each (seed 0), headed by their Panoptic Quality and tumor–CD8 contact F1 over the whole region. **The region:** each method’s whole reconstruction, tumor compartment filled with its 20 *µ*m margin, CD8 cells coloured by where they sit, Tregs as triangles; the dashed square is the 146 *µ*m window magnified below, which holds 14 of the region’s 18 expert contacts. **Cell classes:** every cell by its gated class. **Tumor–CD8 contacts:** the expert’s contacts, black recovered, red dashed lost, blue reported but absent from the reference. At region scale the methods agree; at contact scale they separate: the U-Net and KODIAK are indistinguishable as segmenters here (0.707 against 0.708 PQ, 0.890 against 0.882 segmentation F1) yet keep 5 and 9 of the window’s 14 contacts, and Cellpose, the better segmenter (0.750 PQ), keeps 6. Over all six regions the endpoint orders the first two the other way (0.58 against 0.50; Section 5.5; every region in Table 5).

**Figure 3:**
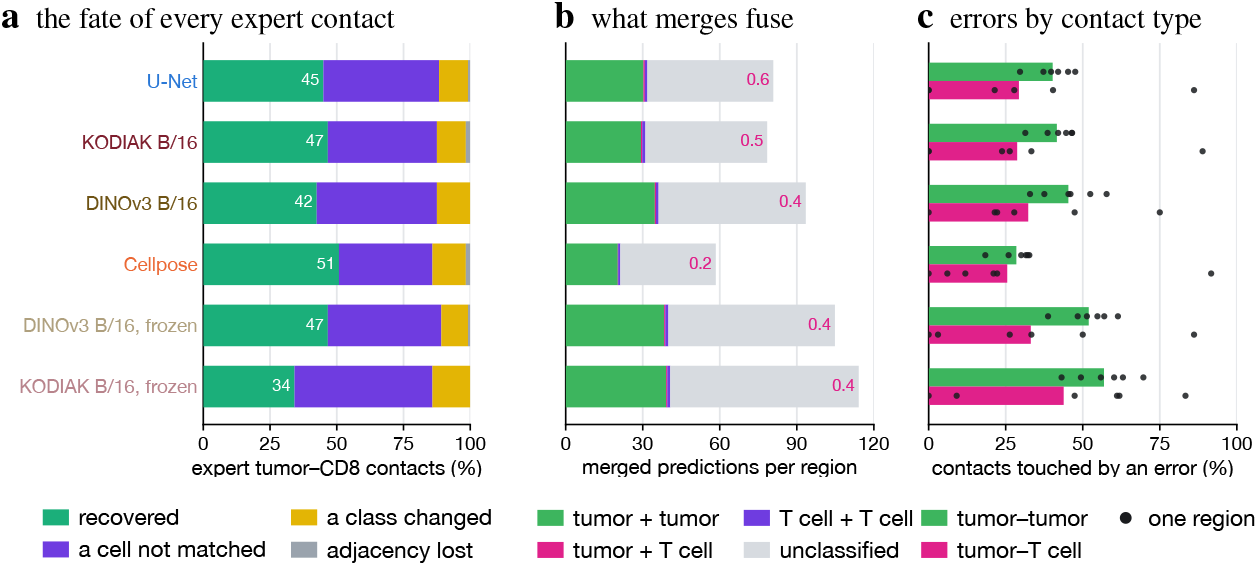
Where the expert’s tumor–CD8 contacts are lost (bars, mean over the three runs). (a) The fate of each of the 40 contacts the expert cells define: *recovered* means both cells matched at IoU *≥* 0.5, both kept their class and the two are still in contact; otherwise a cell was never matched, a class changed, or the adjacency broke (white, the recovered share). (b) Predictions swallowing two or more confirmed cells, by what they fuse; magenta, the heterotypic count. (c) The share of tumor–tumor and tumor–T cell contacts with at least one cell not correctly matched (dots, the six regions).

**Figure 4:**
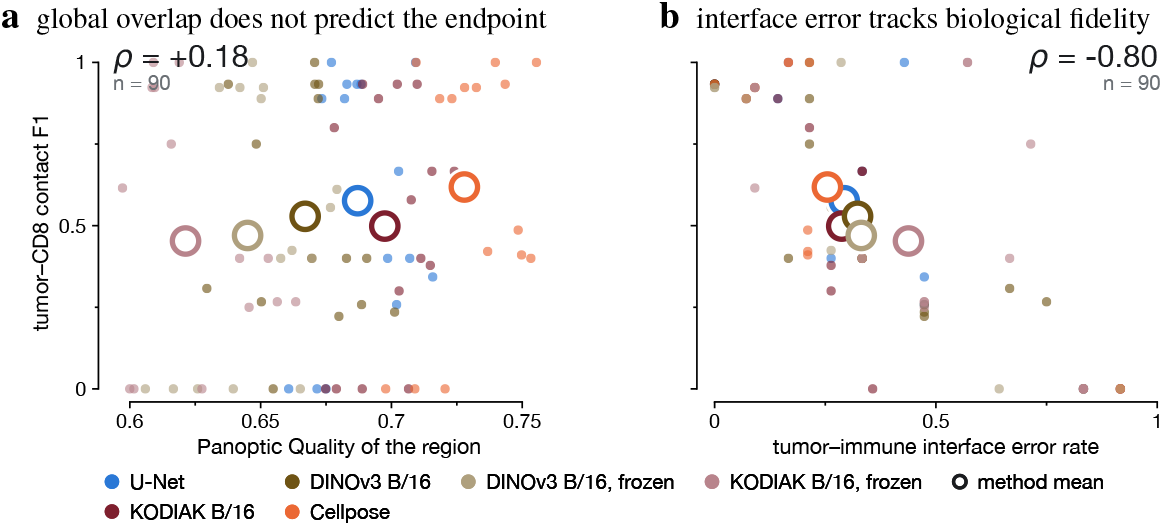
Segmentation quality against biological fidelity, one point per run and region. (a) Panoptic Quality against tumor– CD8 contact fidelity: over the range the methods occupy the two are almost unrelated (*ρ* = −0.18) and the method means (open circles) do not order as PQ does. (b) The same endpoint against the share of expert tumor–T cell interfaces at which a cell is not correctly segmented: this predicts the biological reading (*ρ* = 0.80) where the overlap metrics do not (F1 +0.21, Dice +0.19). The rate counts the endpoint’s own contacts; its independent variant is reported in Section 7. Descriptive scores: Figure 9.

## 6 Discussion

Three lessons emerge. Continued pretraining pays off when the encoder is fine-tuned and not when it is frozen, so its value lies in adaptability rather than decodability, and it closes about a third of the gap to a model pretrained on annotated cells. Every trained model is limited by recall in small, crowded cells. Most importantly the levels do not move together: phenotyping is robust to the segmentation differences we measure, whereas the tumor–CD8 spatial interaction test spreads about twice as widely as segmentation F1 and reorders the methods (Figures 3 and 4). As a focused case study, it exposes a mechanism of propagation. Segmentation errors are not biologically exchangeable: a missed cell at a tumor–immune interface removes an interaction from the inferred ecosystem even when its cost to a global overlap metric is small.

**Table 5.** Tumor–CD8 contact F1 per test region of Slide 2: mean over the three runs, with the range over runs in parentheses. *Contacts* is the number of expert contacts that defines the region’s score; R2 has none, so its score is undefined, and the last row is the mean over the five regions with contacts, the figure reported in Section 5.5.

| Region | Contacts | Fine-tuned |  |  |  | Frozen |  |
| --- | --- | --- | --- | --- | --- | --- | --- |
|  |  | U-Net | KODIAK | DINOv3 | Cellpose | KODIAK | DINOv3 |
| R1 | 18 | 0.33 (0.26–0.40) | 0.42 (0.30–0.58) | 0.24 (0.22–0.26) | 0.44 (0.41–0.49) | 0.26 (0.25–0.27) | 0.53 (0.42–0.61) |
| R2 | 0 | — | — | — | — | — | — |
| R3 | 9 | 0.00 (0.00–0.00) | 0.00 (0.00–0.00) | 0.19 (0.00–0.31) | 0.00 (0.00–0.00) | 0.00 (0.00–0.00) | 0.00 (0.00–0.00) |
| R4 | 4 | 0.93 (0.89–1.00) | 0.56 (0.00–0.89) | 0.88 (0.75–1.00) | 0.93 (0.89–1.00) | 0.92 (0.75–1.00) | 0.63 (0.00–1.00) |
| R5 | 7 | 0.93 (0.93–0.93) | 0.93 (0.93–0.93) | 0.93 (0.93–0.93) | 0.93 (0.92–0.93) | 0.82 (0.62–0.92) | 0.92 (0.92–0.92) |
| R6 | 2 | 0.69 (0.40–1.00) | 0.58 (0.40–0.67) | 0.40 (0.40–0.40) | 0.80 (0.40–1.00) | 0.27 (0.00–0.40) | 0.27 (0.00–0.40) |
| mean | 40 | 0.58 | 0.50 | 0.53 | 0.62 | 0.45 | 0.47 |

The association between interface error and contact fidelity is partly expected, because the metric is evaluated at the interface of interest. The observation that matters is therefore not its magnitude but the failure of global Panoptic Quality, Dice and F1 to distinguish methods that preserve those interfaces differently. For a benchmark meant to support spatial pathology, this argues for reporting at least one biologically targeted endpoint beside the global metrics, and motivates testing whether interface-aware measures transfer across endpoints and tissues.

## 7 Limitations

Our study has several limitations. The benchmark uses three slides of one tissue and one panel in fixed development and test roles; the second test slide replicates the segmentation ordering but is scored at that level only. The reference masks originate from HALO objects followed by expert review, so cells absent from the initial HALO segmentation cannot enter the reference set, and the expert phenotype labels of the test slide have not yet been used to validate the gates there. Phenotype thresholds were calibrated on expert labels from the development slide and may require recalibration across panels, tissues, or acquisition platforms.

The tumor–CD8 analysis is a focused case study of error propagation to a stringent spatial interaction endpoint, not a powered estimate of interaction frequency. Its 40 expert contacts illustrate endpoint sensitivity, complementing the broader phenotype, neighborhood, and niche analyses; they are spread unevenly over six regions, so one region carries none, another only two, and a single missed CD8 cell can remove all of a region’s contacts (Appendix A.5.2). Because interface error is measured where the endpoint is defined, we interpret its association as evidence for a mechanism of propagation rather than a transferable effect size; an independent variant excluding tumor–CD8 interfaces reaches *ρ* =− 0.34. Thus global metrics are insufficient on this dataset for selecting models intended to preserve tumor–immune contacts.

Finally, this study evaluates biological fidelity rather than clinical utility. Larger independent cohorts, additional tissue types and marker panels, and clinically anchored endpoints will be required to determine how broadly these findings generalize.

## 8 Conclusion

Cell segmentation defines the substrate from which phenotypes, neighborhoods and tumor ecosystems are inferred, so how we evaluate it decides which models reach spatial-pathology pipelines. Under a protocol that generalizes to an unseen slide and holds the downstream pipeline fixed, geometric and biological fidelity come apart: phenotype assignment is robust to the segmentation differences we measure, while a stringent tumor–CD8 spatial interaction test spreads about twice as widely as segmentation F1 and reorders the methods. In this focused case study, Panoptic Quality tracks contact fidelity at *ρ* = 0.18, errors localized to tumor–immune interfaces at *ρ* = − 0.80. Errors of similar geometric magnitude therefore carry different biological consequences, and foundation models for spatial pathology should be evaluated not only by how accurately they recover cells, but by whether they preserve the relationships those cells are used to measure.

## A Appendix

Section A.1 states the evaluation hierarchy the paper is organized around. Section A.2 gives the annotation, provenance and composition of the benchmark data, and Section A.3 the architecture, training recipe and prompting settings every method received. Sections A.4 and A.5 hold the per-cell error analysis and the downstream analyses behind Sections 5.1 and 5.5, and Section A.7 lists the code release.

### A.1 The evaluation hierarchy

Table 3 states the four levels the paper reports at and what each asks of a method’s cells. The levels are nested in the sense that each consumes the output of the one below, so an error at the segmentation level can only be amplified or absorbed further up, never repaired. Empirically they compress: the segmentation F1 of the trained rows spans 0.814–0.901, full-type phenotype agreement only 94.3–95.9 %, and whole-graph edge recall 0.669–0.837, while the tumor–CD8 endpoint spreads wider than any of them (0.45–0.62).

### A.2 Benchmark data

Two details of the data that the main text states but does not develop: the human phenotype labels behind the gates, and how much the tissue varies between annotated regions.

#### A.2.1 Expert phenotype labels

The gates of Section 4.3 are calibrated to the labels of Table 4, never to the segmentation reference, and they never enter segmentation training or scoring. The Slide 1 set is a single 1,024^2^-pixel window tagged exhaustively per session, so a cell left untagged is a negative rather than a missing label; that convention is what makes the window usable for calibrating a threshold rather than only for confirming positives. The Slide 2 set, four such windows tagged by the same tool, is held out with the slide and enters no number reported here. The Slide 3 set is bounding boxes over two reviewed fields, which locate cells of a type but never define a boundary.

#### A.2.2 Tissue archetypes

Figure 6 draws the two regions whose depth profiles are flat in Figure 5b. They differ in level rather than in shape: R06 of the development slide is 71 % tumor and 10 % immune, R05 of the test slide 28 % tumor and 44 % immune. Together with the graded region of Figure 5 they bracket what a single annotated region can look like, which is the variation the region-level protocol averages over and the reason we report per-region differences rather than pooled counts.

**Figure 5:**
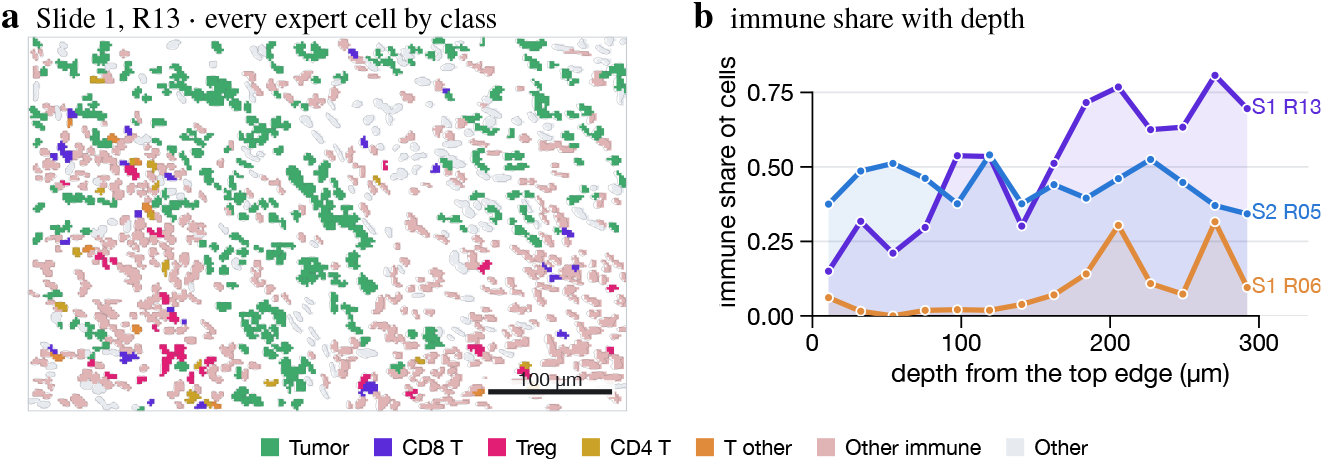
Composition varies within a region, not only between regions. (a) Every expert cell of one annotated region of the development slide by its gated phenotype (Section 4.3), flagged objects removed. (b) The immune share of cells in fourteen horizontal bands, here and in two further regions. R13 is graded, rising from 0.15 at the top edge to 0.80 at the bottom as tumor gives way to a CD45^+^ infiltrate carrying most of its CD8 and Treg cells; a tumor-dominant region of the same slide and an immune-rich one of the test slide are flat and differ in level (Figure 6). A region is therefore not one tissue.

**Figure 6:**
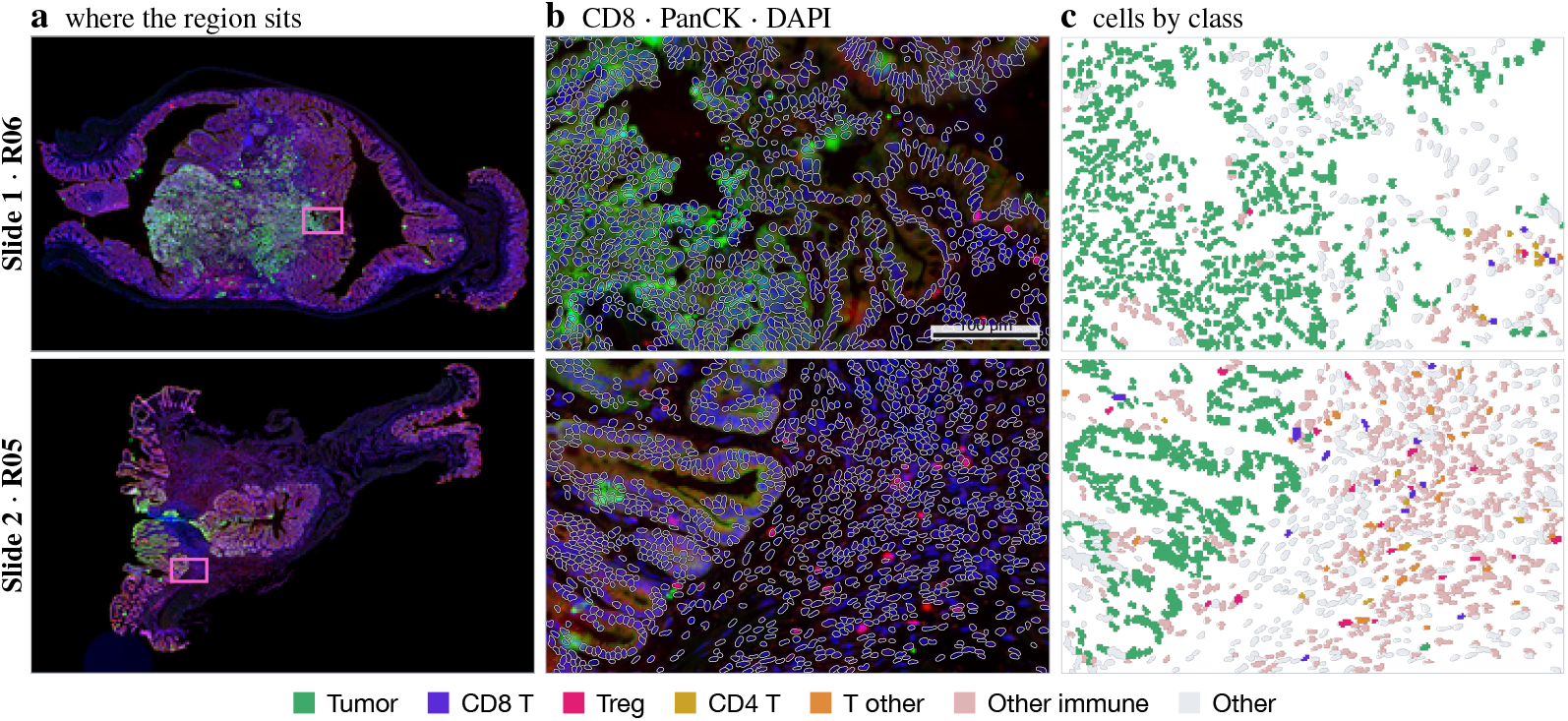
The two flat regions of Figure 5b, each on its own slide. (a) The whole slide in the same three markers as (b), under the same slide-level normalization and curve and at the exposure of Figure 1a–c, so every whole-slide view in the paper matches and the locator and its zoom are one image at two scales; the region is boxed. R06 of the development slide and R05 of the test slide sit in different tissue, which is why they are drawn separately rather than as two crops of one field. (b) The tissue at full resolution, CD8 in red, PanCK in green and DAPI in blue, with every drawn cell outlined individually; the green mass around the box in (a) is the PanCK^+^ epithelium this region sits inside. (c) The same cells in their gated class, under the conventions of Figure 5. R06 is 71 % tumor and 10 % immune, R05 28 % tumor and 44 % immune; both are expert-reviewed and drawn with flagged objects removed.

**Figure 7:**
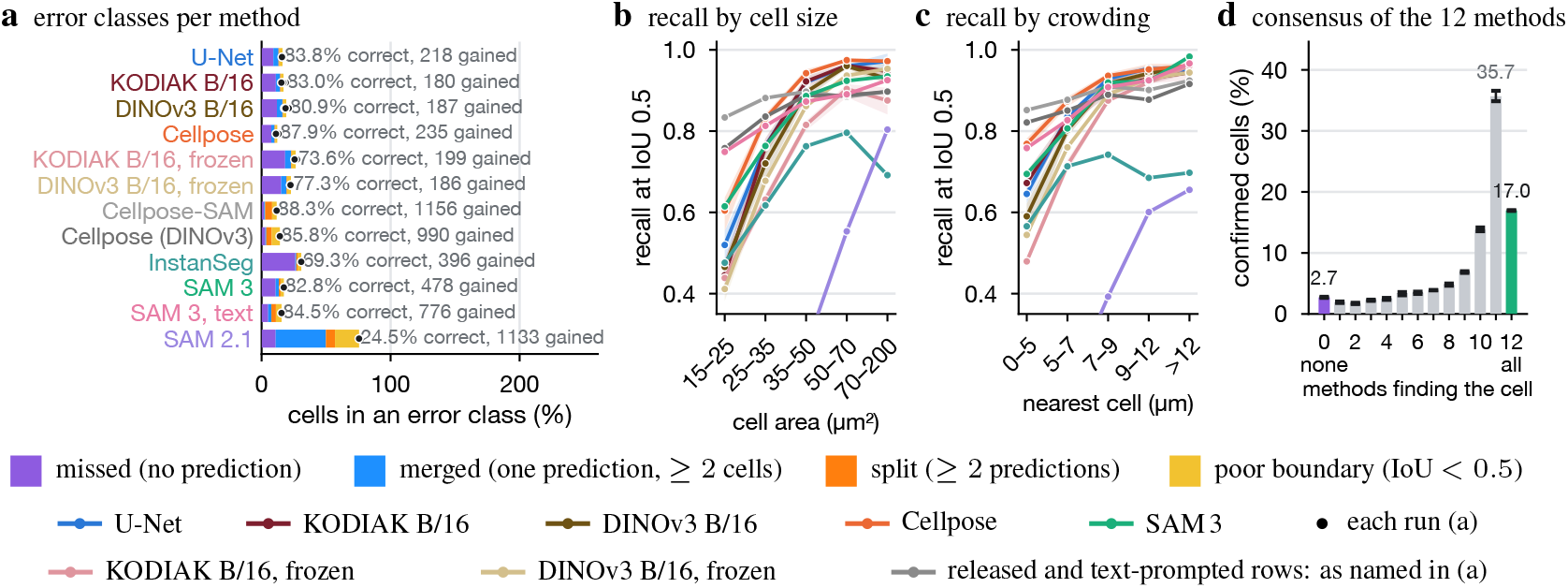
Error analysis on the test slide. (a) Reference cells by error class per method (bars, mean over the three runs; dots, each run; text, the share correct at IoU *≥*0.5 and the gained predictions per run). (b, c) Recall at IoU 0.5 by reference cell area and by distance to the nearest neighboring cell. (d) Share of confirmed cells found by exactly *k* of the 12 methods at matched runs; *k* = 0 are the cells no method finds.

### A.3 Models and training

Every trained row shares one architecture family, one data stream and one optimizer schedule, so that the comparison of Section 5.1 varies only the encoder initialization; the prompted rows receive no gradient updates at all and are specified entirely by their decoding thresholds.

#### A.3.1 Architecture and training recipe

Each marker is scaled to [0, 1] by the 1st and 99.7th intensity percentiles of a fixed sample of tissue tiles of its own slide. The ViT encoder reads the input upsampled 2 *×* (one token per 8 pixels, since cells span 10–20 pixels) under a light UNETR-style [17] decoder over blocks 3, 6, 9 and 12; the frozen variants train only that decoder. The loss is 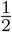 MSE on 5*×* -scaled flows plus binary cross-entropy, masked by the ignore mask, and decoded objects below 15 pixels are discarded. Every model sees the identical stream of 256^2^ tiles with seeded flips and right-angle rotations and no photometric augmentation, for 10 epochs at an effective batch of 16, under AdamW [32] at peak learning rate 2 *×* 10^−3^ with layer-wise decay 0.75 down a pretrained encoder, two warm-up epochs and cosine decay. Every run trains on one Apple M-series GPU: about 5 minutes for the U-Net, 20 minutes for a frozen-encoder row and 40 minutes for a fine-tuned ViT-B/16 row; scoring a run on both held-out slides takes about a minute.

### A.3.2 Prompting settings

SAM 3 with exemplars: detections below score 0.5 are dropped, masks are thresholded at 0.5, instances outside a plausible area range are removed, and overlapping detections are resolved by score with duplicates suppressed above 50 % overlap; the result is stable across exemplar draws. SAM 3 with text: the noun “cell” at detection threshold 0.3, since text-prompt scores run systematically lower than exemplar scores (the phrasings “a cell” and “nucleus” return no detections). SAM 2.1: similarity peaks above 0.7 at a minimum distance of 6 pixels, each decoded from a box prompt of 2.4 *×* the exemplars’ median radius, candidates with predicted IoU below 0.7 discarded, and the same area and duplicate filters.

### A.4 Segmentation error analysis

The summary metrics of Table 2 say how much a method misses but not what it misses. This section defines the per-cell classes behind Figure 7 and reports recall by cell size and crowding, together with the consensus of the twelve methods: 17.0 % of confirmed cells are found by all of them, 2.7 % by none, and 2.3 % by no run of any method, which bounds the recall attainable against this reference.

#### A.4.1 Error taxonomy

A reference cell is *correct* at IoU *≥* 0.5; *missed* when no prediction covers a fifth of it; *merged* when its prediction also covers at least half of another cell; *split* when two or more predictions cover it; *poor boundary* for a single prediction below IoU 0.5. Recall by cell size and crowding and the consensus analysis of the cells no method recovers are in Figure 7.

#### A.4.2 What the methods predict

Table 2 and Figure 7 count the errors; Figure 8 visualizes them. Every reference cell carries the class it was counted as, so the figure and the table describe the same cells. Two things are visible that a table cannot show. The trained models fail on the same cells — small nuclei inside dense epithelial bands — rather than on scattered ones, which is why their recall separates while their precision does not. And Cellpose-SAM, applied without adaptation, recovers more reference cells than several trained models while filling the gaps between them with objects that match no reference cell, the failure its count ratio of 1.26 reports and its F1 of 0.781 hides.

**Figure 8:**
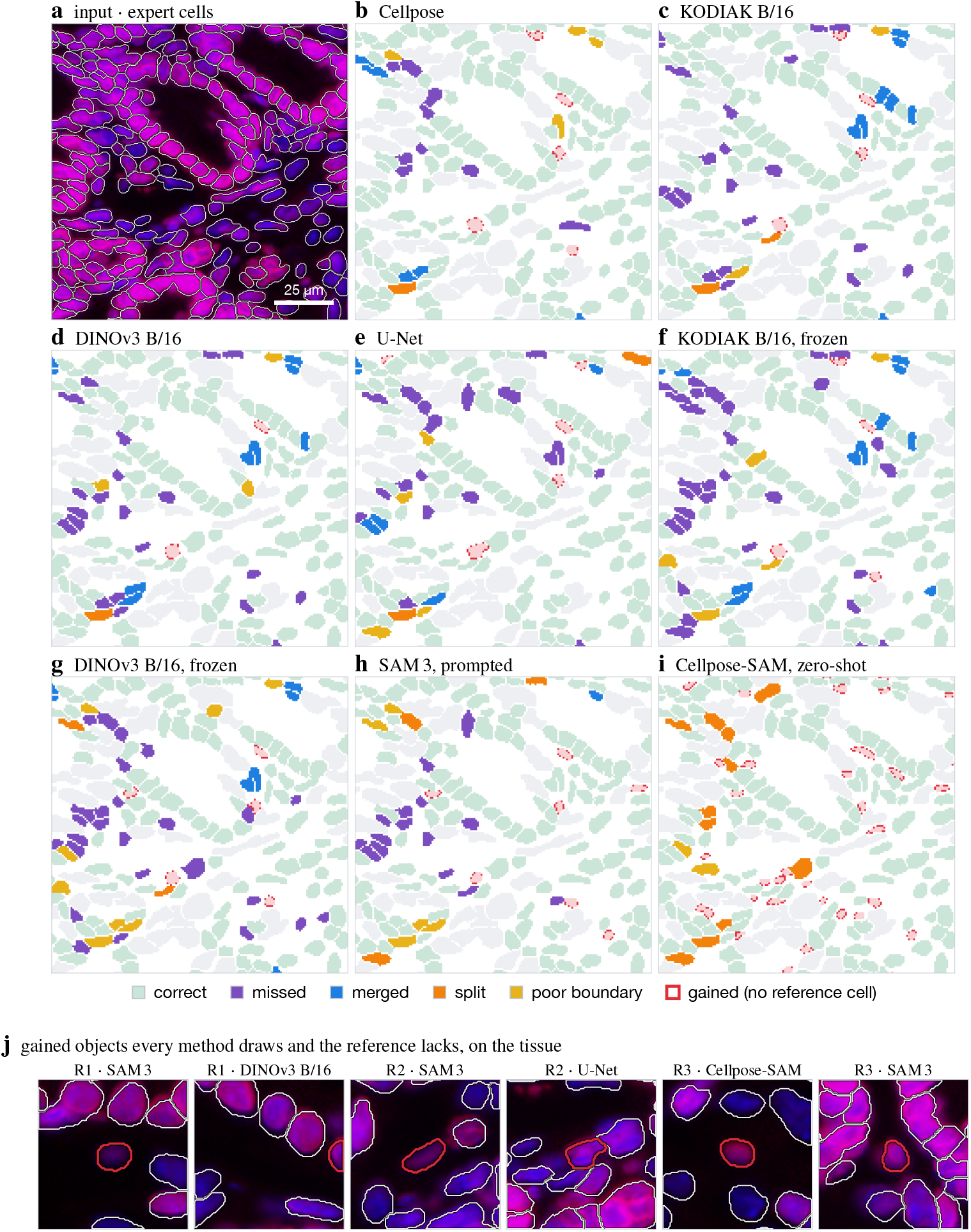
What each method predicts, on the densest 125 *µ*m window of test region R2. (a) The input the two channels form — LAP2 in red and DAPI in blue, green empty, the display blend of Figure 1 — with every confirmed expert cell outlined individually. This is what every method in (b–i) reads; the three-marker view of Figure 6b draws different channels and is not comparable in colour. (b–i) The same cells, each filled with the error class it was counted as in Table 2 and Figure 7: the taxonomy is computed on the whole region under the paper’s protocol and ignore mask, not on this window, so the colours are the counted ones. Cells the expert flagged are ignored everywhere and drawn pale grey; predictions covering no reference cell are filled pink. Seed 0 throughout. (j) Six of those extra objects, each drawn on the tissue at 34 *µ*m with the expert’s cells in white and, in red, the contour drawn by the method each panel names — the object is placed by all eight, and the named one is simply the largest of the eight outlines at that spot.

**Figure 9:**
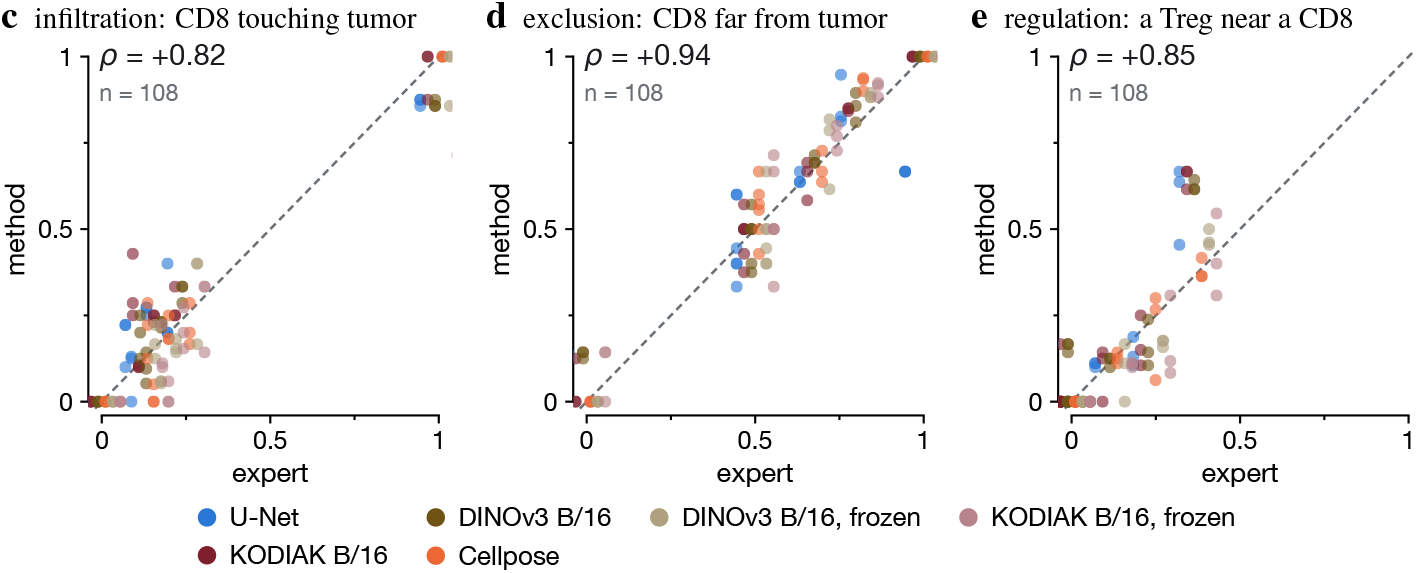
Each method’s descriptive score against the expert’s on the same region, with the identity line: CD8 cells touching tumor, CD8 cells further than 20 *µ*m from the tumor compartment, and CD8 cells with a Treg within 20 *µ*m. Points are offset horizontally by method, by up to *±* 0.055, so the runs sharing an expert value stay separable and a method can be followed across the strip. These three are reproduced closely; the contact endpoint of Figure 4, which needs both cells of a pair to survive, is not.

The last row asks what those extra objects are. Across the six test regions, 560 of them sit where at least two methods independently place a cell and the reference has none, and 70 where all eight do. The six drawn are the largest of the unanimous ones: each is an object every method segments, none of them shares an error mode, and the expert’s own contours around it are drawn in white. Several enclose visible nuclear signal, so part of what the metrics charge as false positives is plausibly annotation the reference is missing rather than error — which is why recall against this reference is bounded (Figure 7d) and why an independently re-annotated field is the one thing that would settle it (Section 7).

### A.5 Downstream analysis

Three companions to Section 5.5: how the region of Figure 2 was chosen, the contact endpoint region by region, and the three descriptive endpoints that the methods do reproduce.

#### A.5.1 How the region of Figure 2 was chosen

The region is chosen by a rule fixed in advance: among the six test regions with |ΔPQ| *<* 0.02 between the U-Net and KODIAK and at least eight expert tumor–CD8 contacts, the one maximizing the difference in contact F1. R1 wins it: over the whole region the two differ by 0.001 in PQ and 0.008 in segmentation F1 but by 0.18 in contact F1 for the runs drawn, and by 0.09 averaged over the three runs of each, across the 18 expert contacts the region contains. The direction there is the reverse of the six-region mean (Section 5.5), which is the point: at matched Panoptic Quality the endpoint can go either way, and per region it does (Table 5).

#### A.5.2 The contact endpoint region by region

Table 5 gives tumor–CD8 contact F1 for every trained row in every test region, as the mean and range over the three runs, beside the number of expert contacts that defines it. Two features limit what any single region can carry. Its denominator is small and the 40 contacts are spread unevenly, so in a region with few contacts a single missed CD8 cell can remove all of them. And the endpoint is less stable across runs than segmentation: at these denominators one run can recover every contact of a region while another recovers none. This is why the main text reports the six-region mean, treats close pairs as unresolved, and draws Figure 2 under a rule fixed in advance rather than by inspection.

#### A.5.3 The three descriptive endpoints

Infiltration, exclusion and regulation are region-level fractions, and every trained method tracks the expert closely on all three (Spearman +0.82, +0.94 and +0.85 over the 108 run *×*region points). They are counted over the cells a method finds, so a missing cell shifts a fraction slightly; the contact endpoint of Figure 4 instead needs both cells of a pair to survive, which is why it separates the methods where these do not.

### A.6 What continued pretraining changes in the representation

The comparison of Section 5.1 varies only the encoder initialization, so what separates those rows is whatever continued pretraining did to the representation. This section reads that representation directly, on the test slide, using the ViT-B/16 checkpoint the paper scores and the DINOv3 weights it was initialized from. Every 16-pixel token of the six test regions is encoded by both, giving 35,511 tokens of which 6,052 (17 %) hold an expert cell and inherit its reference niche.

#### A.6.1 What a token can be read out of

Leave-one-region-out linear probes over the six regions, balanced accuracy; both encoders read the three-channel input of Figure 10a, with PanCK on green, unlike the segmentation input. At full width (768 dimensions for both encoders) the reference niche is recovered at 0.502 from KODIAK tokens against 0.436 from DINOv3 tokens, and cell lineage at 0.656 against 0.559; with both encoders reduced to their first 64 principal components, which limits the probe’s capacity relative to six training regions, the niche probe is level (0.505 against 0.518) and the lineage probe keeps its margin (0.686 against 0.629). Raw marker pixels reach 0.686 for lineage and 0.496 for niche, so the tokens of either encoder add little to the markers on the identity question, and the niche advantage of KODIAK tokens at full width does not survive the reduction, so we treat it as tentative. The division of labour one would expect, markers saying what a cell is and tokens what surrounds it, is consistent with these numbers but not established by them.

**Figure 10:**
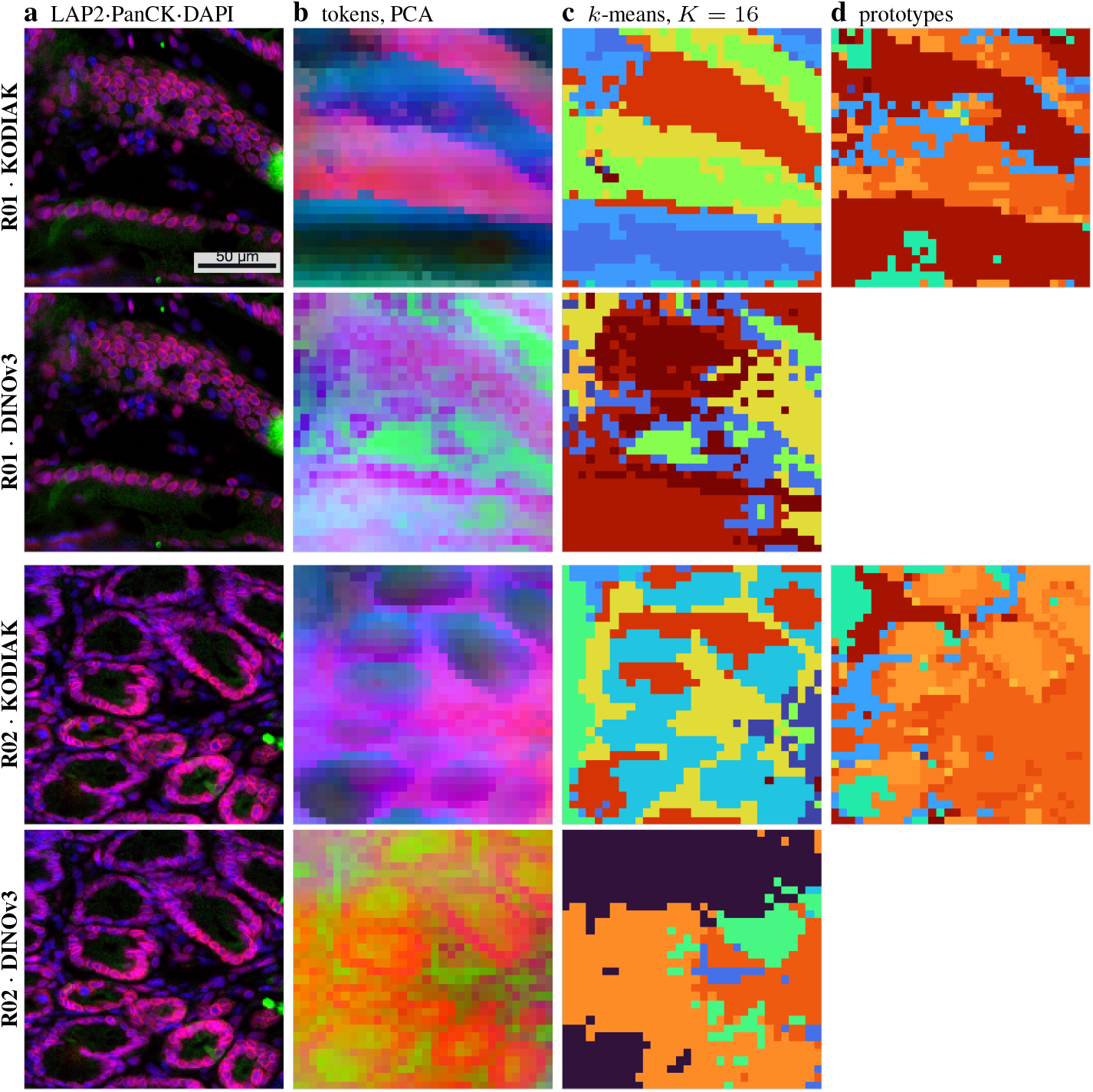
The same tokens through both encoders. Two test regions, each read by KODIAK and by the DINOv3 initialization it continues from, on a 166 *µ*m window chosen as the densest in expert cells. (a) The input both encoders receive. (b) The token embeddings projected to their first three principal components and shown as RGB, fitted per encoder, so colour is similarity of representation and not of marker. (c) The same 768-dimensional tokens partitioned by one *k*-means with *K* = 16, fitted separately but identically for each encoder, so the two rows of a region differ only in the features underneath; this is the matched read-out and not KODIAK’s own prototype codebook. Colours are arbitrary and are not comparable between rows. (d) That codebook: the prototype each token is assigned by KODIAK’s own quantizer, the argmax over its 128 prototypes in the 256-dimensional projector space. It is a learned quantization rather than a matched read-out, and the released DINOv3 checkpoint carries only the backbone — its pretraining head is not distributed with the weights — so the DINOv3 rows have no such map (Section A.6.2). KODIAK’s map is the smoother of the two, with larger contiguous territories that follow the epithelial bands, while DINOv3’s varies at a finer spatial scale.

This does not contradict the frozen rows of Table 2, where KODIAK falls behind its DINOv3 control. Those rows train a dense decoder to delineate cells from the same tokens; the probes here ask what a token says about its neighborhood. A representation can carry more about tissue context and less about the boundary of the cell inside a patch, and the two measurements are not interchangeable.

#### A.6.2 What its clusters correspond to

Quantizing the same tokens with one *k*-means per encoder gives the code maps of Figure 10c, which put the two partitions side by side on the same windows. They differ in character: KODIAK’s territories are larger and follow the epithelial bands, while DINOv3’s change at a finer spatial scale. What a linear probe can decode from a representation and how that representation partitions under clustering are different questions, so we report the probe of Section A.6.1 and these maps separately rather than treating either as evidence for the other.

KODIAK’s own prototype layer is also unevenly used at this checkpoint: 46 of 128 prototypes are live, the most frequent takes 32 % of tokens, and the normalized entropy of the assignment is 0.38 against 1 for uniform. The prototypes live in the 256-dimensional projector space rather than the 768-dimensional token space, so they are reported here as a property of the objective and never compared with the token clusters above.

### A.2 Code and data release

The code is released at https://github.com/marjanstoimchev/beyond-segmentation-accuracy: the data pipeline, every segmenter and its configuration file, the training and evaluation commands, the per-region and per-run metrics of every run behind Table 2, the summary tables of the error-taxonomy, consensus, crowding, phenotype, neighborhood, niche and tumor–immune analyses, and one executed notebook per analysis. The slides, the HALO contours and the expert review are private clinical research data and are not distributed; they are available from the corresponding author under a data-use agreement, and the released pipeline runs unchanged on any CellDIVE and HALO export placed in the documented layout.

## Notes

### Competing Interest Statement

The authors have declared no competing interest.

